# SEEK to Identify Super Enhancer-Expression Correlated Pairs using Single-cell Multi-omic Data

**DOI:** 10.1101/2022.11.07.515509

**Authors:** Guoshuai Cai

## Abstract

Super enhancers (SEs) drive cell identity and disease related genes. However, current methods for studying associations between SE and gene expression are time consuming, costly and with poor scalability.

This study formulated a computational approach for screening genome-wide SE-expression associations by analyzing single-cell multi-omic data of transcriptome and H3K27ac histone modification. A pipeline was also constructed for an easy workflow application. Further our application study identified expression correlated SEs (eSEs) in brain and found they mark cell types. Moreover, our analysis provided new insights into the functional role of SEs close to *Kcnip4* and *Nifb1* in frontal cortex neurons and CGE derived inhibitory neurons, linking to neuron development and neurological diseases.

Collectively, this study provides a new tool for studying SE-expression associations and identifying significant expression associated SEs, which pave the way for understanding the regulatory role of SEs in gene expression and related cellular and disease development.

## Introduction

Super enhancers (SEs) are large clusters of enhancer elements, which have been found to be cell-type specific and activate the expression of cell identity genes. Different with the typical enhancers, SEs bind to high levels of master transcription factors and Mediator coactivators and drives high-level expression of their associated genes [1]. Stimulating oncogenic transcription and other processes, SEs play key roles in the pathogenesis of various diseases, including cancers, neurodegeneration and others [2]. Therefore, understanding the regulatory role of SEs on gene expression will provide new insights into the cell and disease development. For example, human brain is functional as a complex and balanced network of excitatory glutamate neurons and γ-aminobutyric acid-containing (GABAergic) inhibitory interneurons. Most of the GABAergic inhibitory neurons are derived from ganglionic eminences (GEs). Interestingly, three GE subregions: lateral, medial and caudal GEs (LGE, MGE and CGE) generates spatially and functionally distinct interneurons. Although still elusive, the gene expression regulatory programs in these three highly diverse interneurons are considered to play important roles in interneuron development and migration, and underlie the links of disturbance of excitatory/inhibitory balance with neurodevelopmental and neuropsychiatric disorders [3, 4].

Genome-wide SE-expression associations will provide key data for understanding the SE roles in gene expression regulation and screening disease/phenotype related SEs. However, current studies mainly rely on knock-out experiments on a single or a few SEs which are time consuming and costly, and with poor scalability. Recent developed multi-omic platforms enable a multi-modal measurement of genomic features in single cells, which provides an unprecedented opportunity in efficient screening of their associations. For example, Paired-Tag [5] simultaneously profiles single-cell transcriptome and histone modifications, such as H3K27ac histone modification which is a reliable surrogate marker for SE identification [2], and therefore provides a new promise in identifying cell type-specific SEs and evaluating their association with gene expression. In this study, we developed a computational approach, SEEK, to unlock this opportunity in assessing <u>S</u>uper <u>E</u>nhancer-<u>E</u>xpression <u>C</u>orrelations that are specific to particular cell types.

The workflow and pipeline of the proposed approach were designed and constructed for evaluating SE-expression associations and identifying significant expression associated SEs (eSEs) by analyzing single cell multi-omic data of transcriptome and H3K27ac histone modification. In the following application study, eSEs were identified in mouse brain and found to be activated in specific cell types. The synchronized activeness of eSEs and gene expression in neuron subtypes provide new evidence of the important role of SEs in neuron development and neurological diseases.

## Methods

### Datasets

Multi-omic data of H3K27ac modification and transcriptome of single cells from the frontal cortex and hippocampus tissues of Male C57BL/6J mice were generated by Paired-Tag [5]. The processed RNA sequencing read counts were downloaded from GEO GSE152020 and the raw protein binding DNA sequencing reads were downloaded from the SRA project of PRJNA638110.

### Single-cell transcriptomic data processing and cell type identification

Single-cell transcriptomic data were processed and analyzed using Seurat [6]. Briefly, the RNA sequencing read counts were normalized using the “LogNormalize” method and further centered and scaled by standard deviations. 2000 highly variable feature were selected, and dimension of data space was reduced using the top 20 principal components. Further, cell clusters were identified using the Louvain optimization-based clustering method and were visualized in reduced dimensions of Uniform Manifold Approximation and Projection (UMAP). Based on the expression of markers that were used in the original study [5], 18 cell types were identified, including 7 FC frontal cortex neuron types (FC L2/3, FC L4, FC L5, FC PT, FC NP, FC CT, FC L6), 3 hippocampus neuron types (HC CA1, HC CA2/3, HC DG), 3 Inhibitory neurons (InNeu-CGE, InNeu-Sst, InNeu-Pvalb), Oligodendrocyte precursor cells (OPC), Oligodendrocytes (Oligo), Astrocytes, Endothelial cells and Ependymal cells. The cell clusters and corresponding marker expression profile were shown in Figure S1.

### Single-cell histone modification data processing and SE identification

Single-cell protein binding DNA sequencing reads were processed following the method of Zhu et al. [5] for barcode decoding and genome mapping to the mm10 mouse genome assembly. Then, in cells of each major types of neurons (i.e., frontal cortex neurons, hippocampus neurons and Inhibitory neurons) which were identified by above transcriptome analysis, the peaks of H3K27ac signal were identified by MACS2 [7]. From these peaks, SEs were identified by ROSE [1] for each cell type, and further sequencing reads of H3K27ac binding DNA at SE regions were counted for quantification in each cell.

### eSE identification

In a particular cell, the signals were normalized by the size of all reads with a consistent scale factor, 10000, and log transformation was applied to transform data to approximately conform to normality. In a particular type of neuron, the correlation between each SE-expression pair will be assessed and used to identify eSEs. However, single cell sequencing data is with high sparsity due to the high drop-out rate, substantial technical noises and biological heterogeneity (e.g., the variation in transcriptional bursting [8]). Typical correlation measures such as Pearson or Spearman’s correlation coefficients can’t provide a robust estimation of correlations from single cell sequencing data. To address this issue, a weighted approach is formulated in this study. Specifically, by adapting a mixture model used in [9], the single cells sequencing count *Y*_*j*_ for the *j* − *th* gene is modeled by a zero-inflated negative binomial (ZINB) mixture model with two components, that

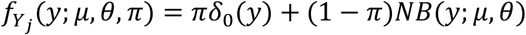

Where the NB distribution with shape (*μ*) and rate *θ* parameters models observations from actual biological material and the Dirac delta distribution *δ*_0_(*y*) models an observed zero from drop-out (i.e., an excess zero). *π* is the drop-out rate. Parameters (*π, μ and θ*) were estimated by fitting the model with a penalized maximum likelihood estimation procedure [9]. By Bayes’ rule, the posterior probability of an observation from actual biological material can be obtained as

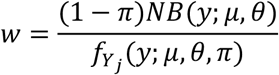

Obtained from scRNA-seq and Paired-Tag data respectively, the posterior probability *w*_*e*_ and *w*_*e*_ were further considered as weights to obtain a robust estimation of correlations based on the weighted ranks of values of expression (*R*_*e*_) and histone modification (*R*_*h*_), that

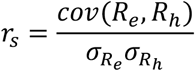

Where,

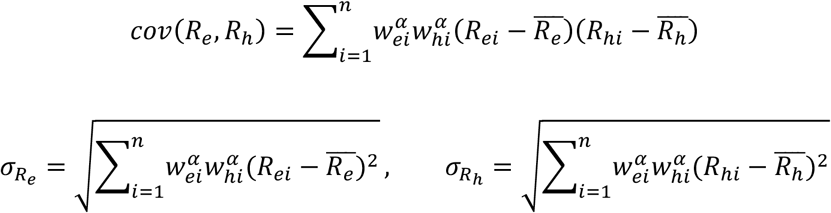

The ranks *R*_*e*_, *R*_*h*_ can be obtained from weights by comparing the being-ranked value *ε* to each of other values *ε*_*i*_, that

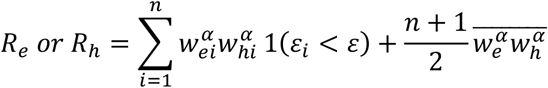

The turning parameter *α* was set to 3 and the wCorr package [10] was used in SEEK to calculate this weighted correlation. Although the probability of that zero is from non-expression but not drop-out is considered, but the correlation calculation is still strongly affected by the number and proportion of missing values. Therefore, features with large numbers of missing values were filtered to achieve a reliable correlation assessment; SE-gene pairs that have non-missing multi-modal data in more than 300 cells and more than 20 cells in each major neuron type were analyzed in this study. eSEs were considered to have a notable correlation with *r* > 0.2 in a particular cell type.

### Cell-type Marker detection and differential analysis

To evaluate if the identified eSEs in this study marker cell types, Wilcox rank sum test implemented in “Seurat” was used to compare gene expression and H3K27ac modification in a certain cell type with that in all other cell types. Bonferroni method was applied to adjust *p* values to control false positives due to multiple comparisons. A cell-type marker was detected with adjusted *p* value < 0.05, log fold change < 0.1, and expression was detected in > 10% of cells. The same analysis strategy was also used to test the difference of *Nfib* gene expression and SE histone modification between InNeu-CGE and the other two inhibitory neuron subtypes.

## Results

### A workflow/pipeline for screening expression associated SEs

To enable the eSE screening, a workflow was designed as illustrated in Figure 1. It is a multi-step procedure for data processing, sequentially with demultiplexing, transcriptome/genome mapping, gene expression quantification, clustering and cell type identification based on single-cell transcriptome data, histone modification peak calling, SE identification, histone modification quantification, multi-modal data conjugation, and eSE identification. The details and tool implementation were described in Methods. The outputs of this workflow are cell type-specific super enhancer regions, single-cell gene expression and histone modification levels, SE-expression correlation coefficients, and identified eSEs. This new approach leverages single-cell multi-omic data of three key advantages: (1) the same-cell joint profiling of transcriptome and histone modification captures cell-to-cell variation which was ignored by traditional bulk data, (2) the large scale of single-cell measurements provides favorable statistical power in association assessment, which was a critical challenge for traditional bulk tissue experiments, and (3) the single-cell resolution provides an opportunity to build SE atlas which has been known to be cell type-specific. In support of the workflow application, a pipeline was constructed, which protocol and codes are available in GitHub (https://github.com/thecailab/KEEP).

**Figure 1.**
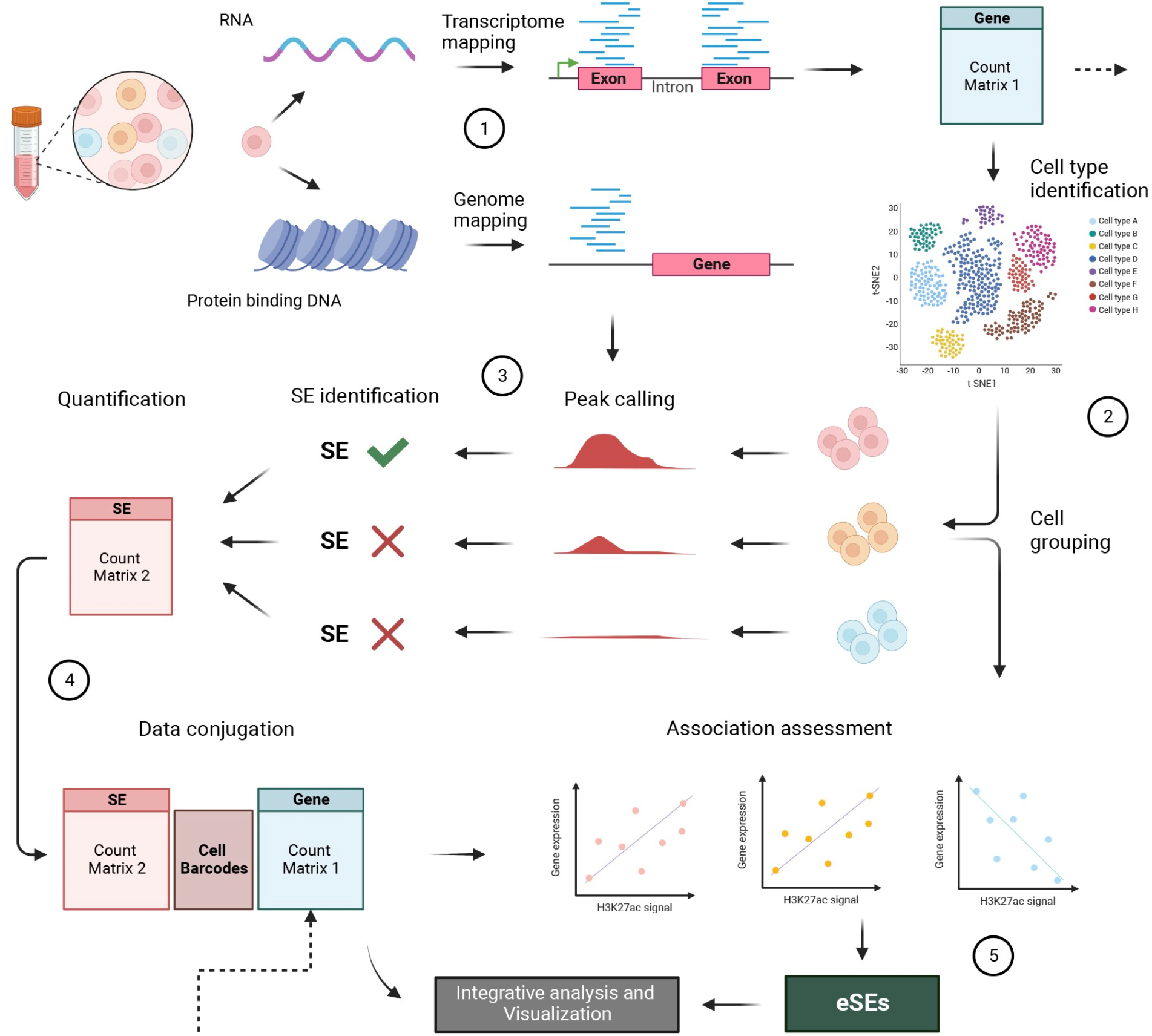
The workflow of EqSE identification using single-cell multiomic data. It constructs a multi-step processing of (1) sequencing data processing, including demultiplexing, transcriptome/genome mapping and gene expression quantification, (2) cell type identification based on single-cell transcriptomics, (3) SE identification and quantification based on cell type-level histone modification data, (4) multi-modal data conjugation, and (5) EqSE identification.

### eSE activeness is cell type-specific in brain

Using SEEK, this study assessed the SE-expression correlation in three major neuron types, frontal cortex (FC) and hippocampus (HC) neurons, and inhibitory neurons (InNeu). As expected, specific SEs were identified these three neuron types (Fig. 2A). Consistent SE-expression correlations were widely found cross these neuron types (Fig. 2B). KEEP identified 7 eSEs, 6 of which showed statistically higher level of histone modification in specific cell types, and significant cell-specific expression were also found in 6 of their associated genes (Fig. 3). As shown in Figure 3 Left, cell-specific expression was found on *Kcnip4* in FC L2/3-6; *Nfib* in FC PT and CT; *Bcl11a, Msra* and *Nfib* in HC cells; *Nfib* in InNeu-CGE; *Eef2k* in FC L2/3. Higher average expression and detection rates were also found on *Phactr3* in OPC, Oligodendrocytes and Astrocytes; *Klf9* in Endothelial cells and *Eef2k* in Ependymal cells, although the differences are not statistically significant which is likely due to the small sample size. Despites of low coverage, histone modification at these eSEs showed a corresponding pattern of activeness according to detection rate and average expression (Fig. 3 Right).

**Figure 2.**
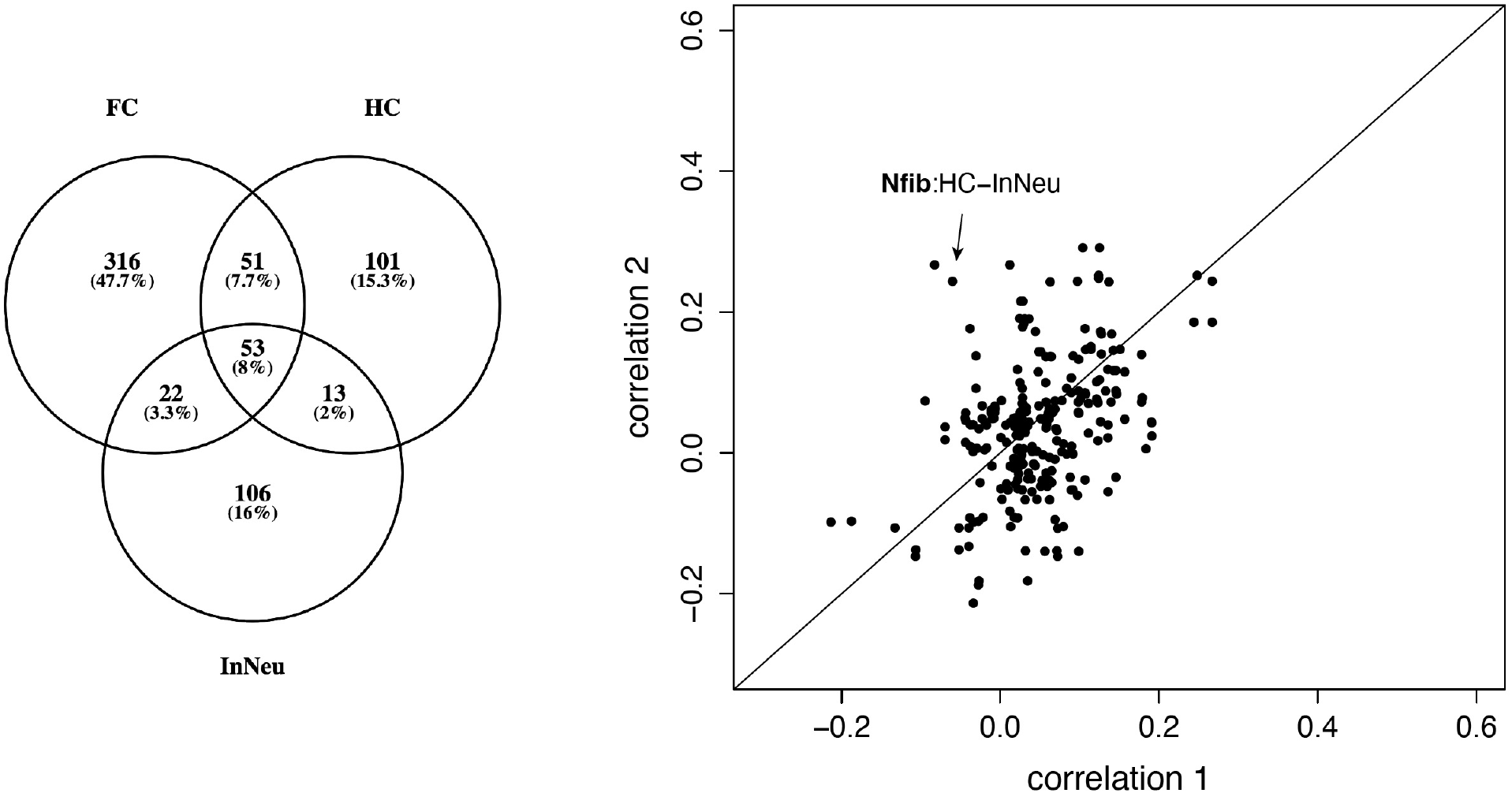
Comparison of SE identification and SE-expression associations in Neuron types. The comparison of SE identification is shown on the left and SE-expression association is shown on the right. For each gene, correlation 1 and 2 are from FC, HC or InNeu for pairwise comparisons. The significant difference of SE-expression associations in *Nfib* between HC and InNeu was identified. FC: frontal cortex; HC: hippocampus neurons; and InNeu: inhibitory neurons.

**Figure 3.**
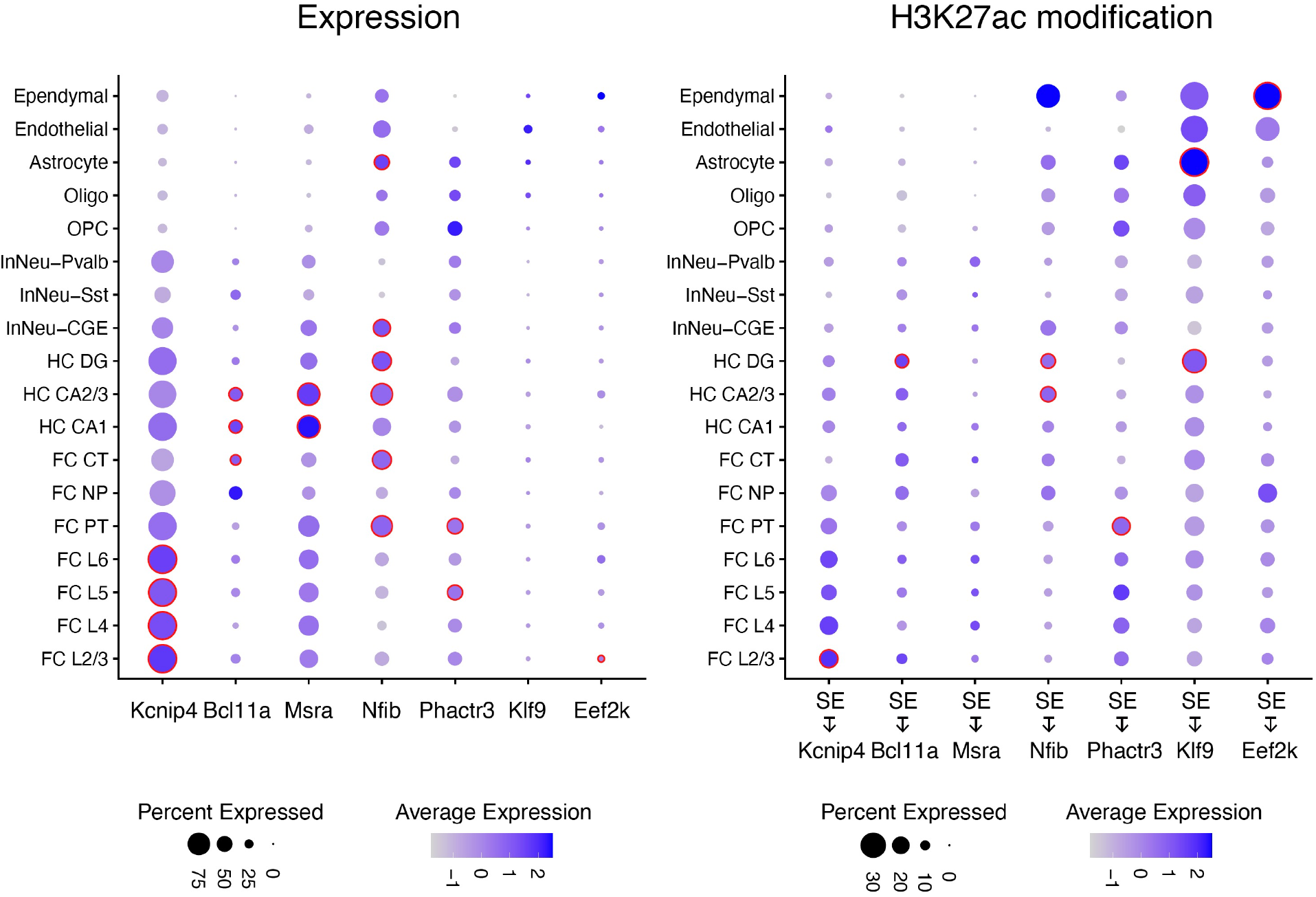
Profiles of eSE features in brain cell types. In a particular cell type, the level of expression of genes (left) and histone modification at each mapped eSE (right) were shown by dot size and color. A larger dot size indicates a higher detection rate of gene expression within each cell type, while a darker color indicates a higher average level of signal. The red circles indicate significantly higher expression in a cell type compared to all others. FC: frontal cortex; HC: hippocampus neurons; InNeu: inhibitory neurons; PT: pyramidal tract excitory neurons; CT: corticothalamic excitatory neurons; NP: near projecting excitatory neurons; OPC: Oligodendrocyte precursor cells, Oligo: Oligodendrocytes

Interestingly, *Nfib* SE-expression association was observed in inhibitory neurons but not in excitatory neurons (Fig. 4B-D), and the association was masked in the pool of all neurons (Fig.4A). As compared to subtypes of MGE derived inhibitory neurons (InNeu-Sst, InNeu-Pvalb), the CGE derived inhibitory neurons (InNeu-CGE) showed synchronized activeness of histone modification at SE and gene expression of *Nfib* with higher detection rate and average level (Fig.4E,F, *p* -value= 1.30 × 10^−32^ for gene expression, *p* -value= 4.99 × 10^−4^ for histone modification).

**Figure 4.**
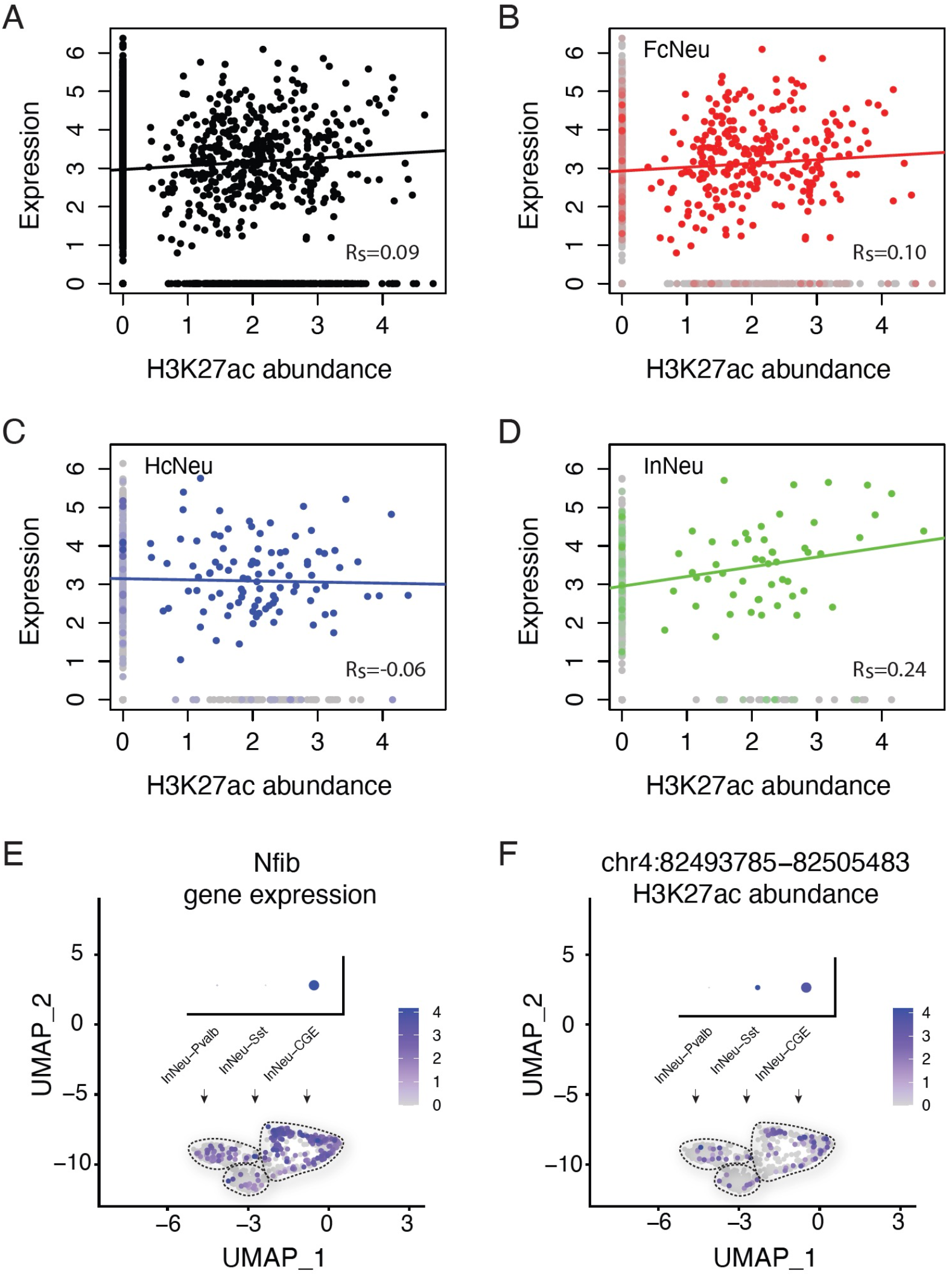
*Nfib* expression and histone modification at proximal SE in neuron subtypes. The top panel shows the SE-expression correlation of *Nfib* in (A) all neuron cells, (B) FC, (C) HC and (D) InNeu specifically. The correlation coefficients were calculated by the weighted methods proposed in this study; the lines were fitted by weighted linear regression. The bottom panel shows UMAP plots of single-cell (E) gene expression of *Nifb* and (F) histone modification at its proximal SE in three inhibitory neuron subtypes. Overlaid dot plots show average detection rates (corresponding to dot size, larger dot indicates higher detection rate) and average signal levels (corresponding to dot color, darker indicates higher signal level). FC: frontal cortex; HC: hippocampus neurons; InNeu: inhibitory neurons.

## Discussion

This study developed a novel workflow with pipeline to study SE-expression association and identify eSEs by interactively analyzing the same-cell transcriptome and histone modification in cell populations. Its performance was cross validated by the broad consistence of associations in three major neuron cell types.

SEs are known to be cell type-specific and thus require methods for cell type-specific analysis. Their cell type-specific patterns could be masked in cell mixtures which are typically used in bulk experiments, as shown in this study that, the analysis of all neuron cells failed to detect the inhibitory neuron-specific SE-expression association of *Nfib*. This study identified eSEs in mouse brain and found their activeness in histone modification and gene expression mark cell subtypes. Of particular interest, a broad SE associated active expression of *Kcnip4* were found in excitatory neurons, especially cortical FC neurons cross L2/3 to L6. *Kcnip4* is known to encode a family member of calcium binding proteins which interacts with voltage-gated potassium channel subunit Kv4 family and regulates transient “A-type” currents (*I*_A_) in neurons and consequently neuronal excitability. [11] Here, our finding provides new insights into the underlying mechanisms of this neuronal excitability regulation, in which SEs may play an active role. Among inhibitory neuron subtypes, the SE and gene expression activeness of *Nfib* was specifically observed in CGE derived inhibitory neurons, indicating SE-involved transcription regulation may contribute to the development and maintenance of inhibitory neuron cell subtypes. Our data is well aligned with an independent report that *Nfib* specifically regulated CGE development [3]. *Nfib* is a member of the phylogenetically conserved nuclear factor I (NFI) gene family which are essential for the development of a number of organ systems. Studies reported that Nfib is important for neural progenitor cell differentiation during cortical development [12, 13]. In this study, the newly identified cell type-specific activation of eSE approximal to *Nfib* may provide a new direction to elucidate its roles in brain development.

A limitation resides in the low per cell sequencing coverage compromises the sensitivity of the single-cell profiles, especially for histone modification detection. This limitation will be remediated by the reducing cost of sequencing and single cell library construction. With necessary modifications, the proposed workflow is versatile and is logically adaptable to other established single-cell multi-omic measures of transcriptome and histone modification, including CoTECH [14], SET-seq [15] and EpiDamID [16], although currently no H3K27ac data from these platforms are publicly available. It is also expected to be easily extended for data from future improved platforms. In addition, the proposed weighted strategy for single cell sequencing data correlation analysis is theoretically applicable to data from other multi-omic platforms to assess the associations between other genomic features (e.g., methylation and gene expression) and inferring gene co-expression networks. We envision a future version of SEEK with this added capability.

## Figures

**Figure S1.**
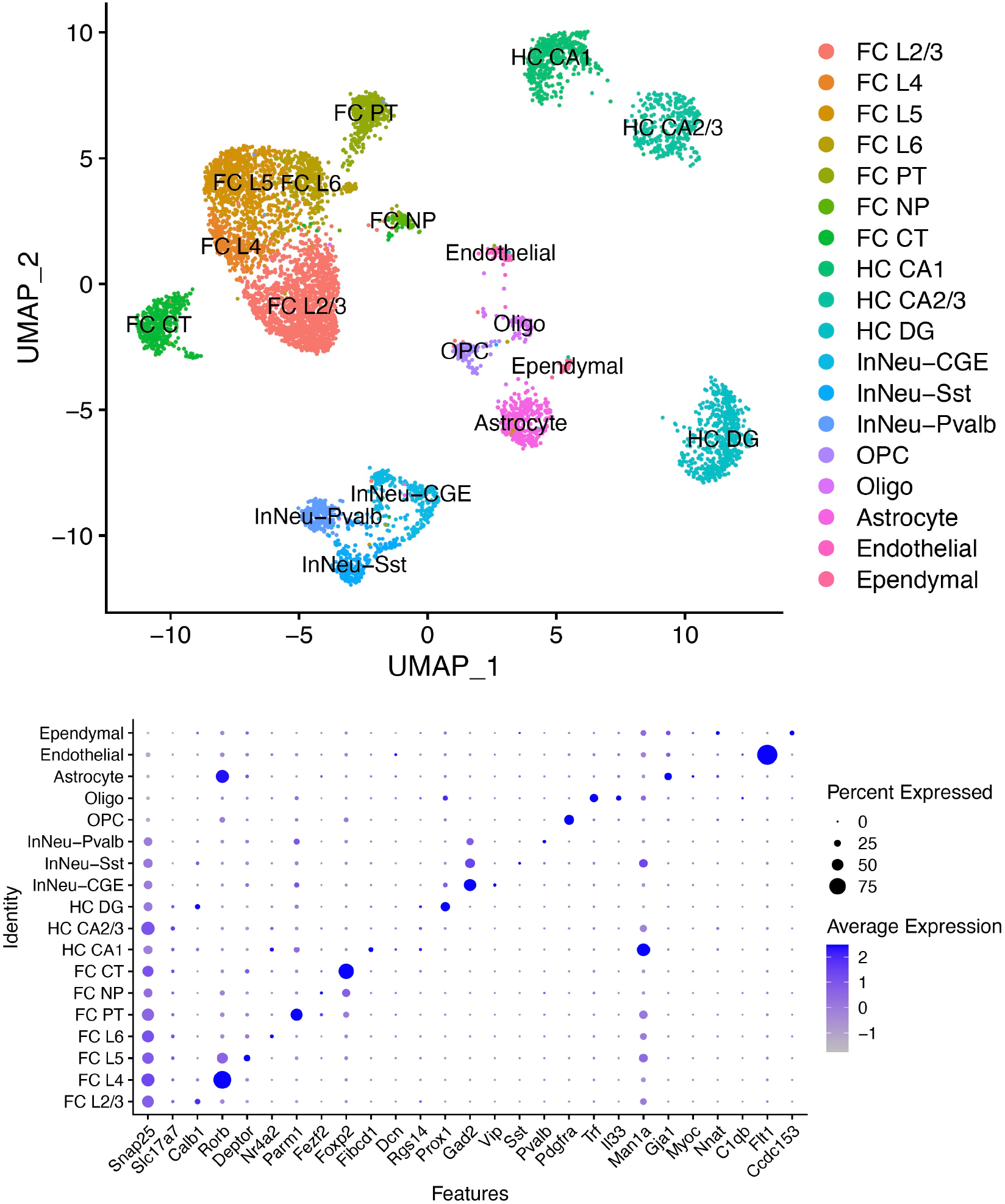
Brain cell clusters and marker expression profiles. For each cell type shown in the UMAP plot (top), the detection rate (corresponding to dot size, larger dot indicates higher detection rate) and average expression level (corresponding to dot color, darker indicates higher expression level) of each cell-type marker are shown in the dot plot (bottom). FC: frontal cortex; HC: hippocampus neurons; InNeu: inhibitory neurons; PT: pyramidal tract excitory neurons; CT: corticothalamic excitatory neurons; NP: near projecting excitatory neurons; OPC: Oligodendrocyte precursor cells, Oligo: Oligodendrocytes

